# Characterization of six environmental *coli*-phages isolated in Astana, Kazakhstan, during the School of Molecular and Theoretical Biology

**DOI:** 10.64898/2026.04.17.719196

**Authors:** Artyom A. Egorov, Konstanty Keda, Oleg K. Klementiev, Jonas Juozapaitis, Daria Akopova, Danya Basalaev, Yuliya Malinouskaya, Uliana Shurlakova, Lisa Trefilova, Aigerim Turgimbayeva, Daria Garshina, Lisa Dialektova, Anna Smolnikova, Maria Markidonova, Julian J. Duque-Pedraza, Polina Selkova, Anu Tyagi, Sailau Abeldenov, Marcus J.O. Johansson, Gemma C. Atkinson, Vasili Hauryliuk, Ilya Terenin

## Abstract

Bacteriophage (phage) collections are essential resources for studying virus–host interactions in bacterial species. Here, we report six *Escherichia coli*-infecting phages that expand the Lund Collection of Bacteriophages. These phages were isolated in 2025 within the framework of the *School of Molecular and Theoretical Biology* for high-school students, from samples collected in Lake Taldykol, Astana, Kazakhstan, using *E. coli* strains MG1655Δ*RM* and EV36 as hosts. The isolated phages comprise Taldykol (LuPh6), a member of the genus *Kagunavirus*; Aidakhar (LuPh7) of the genus *Phapecoctavirus*; Samruk (LuPh8) of the genus *Tequintavirus*; the T-odd-like phage Baiterek (LuPh9) of the genus *Vequintavirus*; and two T-even-like phages Tulpar (LuPh10) and Shurale (LuPh11) that belong to the *Tequatrovirus* genus. This expanded phage collection enhances the toolkit for investigating phage-host interactions and their molecular mechanisms and highlights the use of phage isolation as a component of high school research education.

**Importance:** Phage collections are a key resource for studying phage biology, phage-bacteria interactions and bacterial immune systems. Here, we extend the Lund Phage Collection through the isolation and characterisation of six *E. coli*-infecting phages, including three novel species (LuPh6, LuPh8 and LuPh11) as well as a member of the genus *Phapecoctavirus* that not represented in widely used collections such as BASEL (LuPh7). This study expands the resources available for probing phage-host interactions and demonstrates an example of integrating phage research into education of high school students.

## Introduction

Bacteriophages (or simply phages) are viruses that infect bacteria (Dion *et al*, 2020). Phage collections are an essential tool for research on phage-host interactions. One of the most widely used coliphage collections is BActeriophage SElection for your Laboratory (BASEL), a set of *E. coli* K12-infecting phages isolated by the Alexander Harms lab (Maffei *et al*, 2021), which was recently extended with new genera and O-antigen-dependent phages (Humolli *et al*, 2025). Given the relatively straightforward nature of the phage isolation experiment, many phages have been isolated during educational courses, such as the “phage hunt” organised by the Science Education Alliance (SEA) – a large-scale project in which more than 50 thousands students have isolated in excess of 23K phages (Heller *et al*, 2024). In 2023 our lab at Lund University started building a coliphage collection in the framework of the *Fundamentals of Basic and Applied Phage Biology* course for PhD students and postdocs. The first iteration of the course has yielded five phages LuPh1-LuPh5 isolated on motile *E. coli* strain BW25113 *uspC-flhDC*::IS5 (Shyrokova *et al*, 2025).

Here, we present the results of the second pedagogical phage-hunt project of our lab, conducted during the 2025 *Summer School of Molecular and Theoretical Biology* (SMTB) (molbioschool.org) SMTB 2025 was held from 24 June to 26 July 2025 in Astana, Kazakhstan, and was aimed for school students from around the world. In this project we extend the Lund Collection of phages (Shyrokova *et al*., 2025) by isolating coliphages from Malyi (Small) Taldykol (Kazakh: Кіші Талдыкөл), an urban lake in Astana, Kazakhstan (51°07’00.7”N 71°23’13.5”E). We characterise six environmental *E. coli* phages isolated during the research school and provide their annotation, taxonomic classification, morphology and lytic activity.

## Results and Discussion

To increase the diversity of bacteriophages, we used three *E. coli* strains as the host: i) K-12 MG1655 ΔRM expressing *wbbL* from pAS001 plasmid (Humolli *et al*., 2025), ii) motile K-12 BW25113 *uspC-flhDC*::IS5 (hereafter referred to as VHB17) strain that we used previously to create Lund Phage Collection 1.0 (Shyrokova *et al*., 2025), and, finally, iii) K-12/K1 hybrid EV36 (Vimr & Troy, 1985). Laboratory *E. coli* K-12 strains typically lack O-antigen polysaccharides, and expression of *wbbL* from pAS001 partially restores O-antigen, enabling isolation of phages that depend on it for infection (Humolli *et al*., 2025). BW25113 *uspC-flhDC*::IS5 has upregulated expression of flagella and is well-suited to isolation of flagellotropic phages (Shyrokova *et al*., 2025). The EV36 strain is *kps*^+^, i.e., it expresses the K1 antigen, thus allowing for isolation of phages that use the polysialic acid-containing K1 capsule as a receptor. The K1 capsule is associated with extraintestinal pathogenic *E. coli* (ExPEC) phylogroups that cause invasive extraintestinal diseases (Arredondo-Alonso *et al*, 2023; Cross *et al*, 1984). As K1 can serve as a phage receptor (Gong *et al*, 2021; King *et al*, 2007; Scholl & Merril, 2005), it is tempting to speculate that K1-specific phages could hold promise for anti-ExPEC phage therapy. Two of the strains used lack specific anti-phage defences: BW25113 carries an inactivating mutation in the type I restriction-modification (R-M) system EcoKI (Datsenko & Wanner, 2000; Grenier *et al*, 2014) while MG1655 ΔRM lacks EcoKI as well as McrA, Mrr, and McrBC type IV restriction systems (Maffei *et al*., 2021).

To source the phages, we collected water samples from the Malyi (Small) Taldykol lake. Samples were cleared of debris, then precipitated by zinc chloride, and assayed using the classical agar overlay protocol (Kropinski *et al*, 2009) using the three *E. coli* strains as hosts. Phages were isolated on MG1655 ΔRM strain bearing pAS001 plasmid (ten isolates), EV36 (seven), and BW25113 *uspC-flhDC*::IS5 (VHB17) (one) strain, and their DNA was extracted for sequencing. DNA extraction repeatedly failed for the VHB17-isolated phage, while the remaining seventeen DNA samples were sequenced using Nanopore technology. After deduplication of assembled genomes, six distinct phages were left. The phages were assigned Lund Phage Collection designations (LuPh): LuPh6 and LuPh8-LuPh11 were isolated on MG1655 ΔRM with partially restored O-antigen, and LuPh7 was isolated on EV36. The phages were also given names: one named after the Taldykol lake (LuPh6) and the rest of the names were borrowed from Kazakh mythology: Aidakhar (LuPh7), Samruk (LuPh8), Baiterek (LuPh9), Tulpar (LuPh10) and Shurale (LuPh11).

To classify the novel phages, we first searched for homologous known phages against the NCBI NR database (version of 15 Jan 2025) using BLASTn (Johnson *et al*, 2008) (**Supplementary Table 1**). Genomes for phages related to our LuPh phages were downloaded, and pairwise nucleotide identities were calculated to identify species- and genus-level clusters according to the International Committee on Taxonomy of Viruses (ICTV) criteria (Turner *et al*, 2021; Turner *et al*, 2023). Phages with full-genome nucleotide identity higher than 70% were considered to belong to the same genus, whereas those with an identity above 95% were classified as the same species. We identified three novel species – LuPh6, LuPh8 and LuPh11 – with LuPh6 being the most divergent, sharing only 74.2% nucleotide identity with the closest described virus, *Escherichia* phage vB_EcoS_UTEC10 (Rajab *et al*, 2024) (**Table 1**, **Supplementary Table 2**). Proteome-level genomic alignments of the novel LuPh phages and their top ten closest relatives by nucleotide similarity as listed in NCBI as of 15 January, 2026, are shown on **Supplementary File 3**. The following genera were assigned to the LuPh phages: LuPh6: *Kagunavirus*; LuPh7: *Phapecoctavirus*; LuPh8: *Tequintavirus*; LuPh9: *Vequintavirus*; and LuPh10 and LuPh11: *Tequatrovirus* (**Table 1**).

**Table 1.**
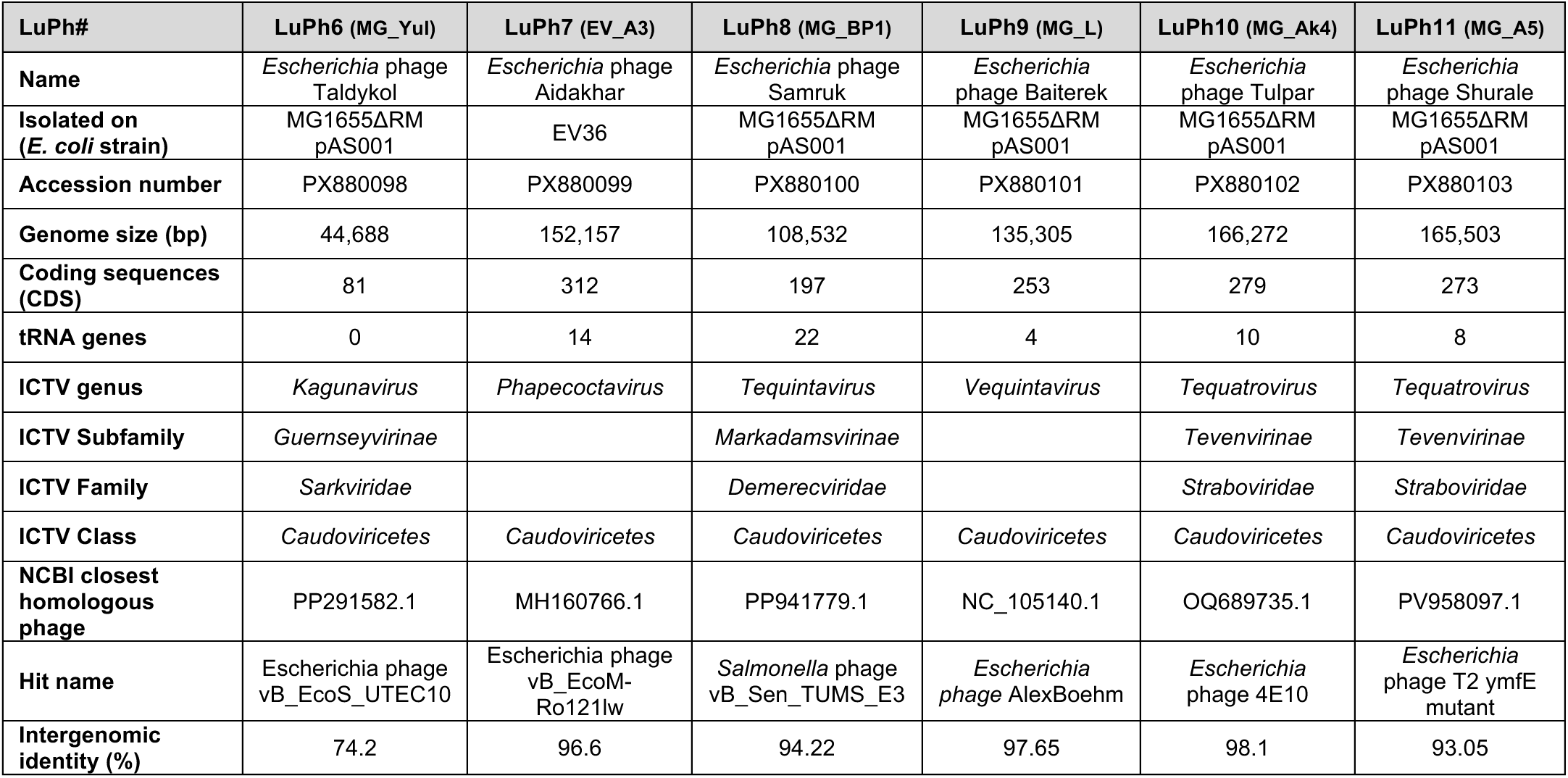
Phages isolated in this study.

To determine how the newly isolated phages expand LuPh 1.0, BASEL and Durham coliphage collections that we use in our lab (Humolli *et al*., 2025; Kelly *et al*, 2023; Maffei *et al*., 2021; Shyrokova *et al*., 2025), we have annotated the assembled genomes using Pharokka (Bouras *et al*, 2023) and then used LoVis4u (Egorov & Atkinson, 2025) to visualise the genome organisation (**Figure 1A**) as well as to calculate proteome composition similarity across the phage set. Full-scale LoVis4u genome alignments of our six phages and those from LuPh 1.0, BASEL and Durham collections can be found in **Supplementary File 4**. LuPh6, a member of the genus *Kagunavirus*, shares the same genus as the Bas70-Bas76 phages from the recently expanded BASEL collection (Humolli *et al*., 2025), sharing ∼69% nucleotide identity with its closest collection relative, Bas73. LuPh7 does not have close representatives in the collections listed above. However, its core genome organisation is similar to that of Bas60-Bas62 (genus *Phapecoctavirus*), with nearly half of protein clusters shared (**Figure 1**; **Supplementary File 4**). LuPh8 belongs to *Tequintavirus* genus (T5-like) and LuPh9 is a member of the *Vequintavirus* Bas48-Bas59 cluster. Finally, LuPh10 and LuPh11 represent *Tequatrovirus* genus (T-even) (**Figure 1B**).

**Figure 1.**
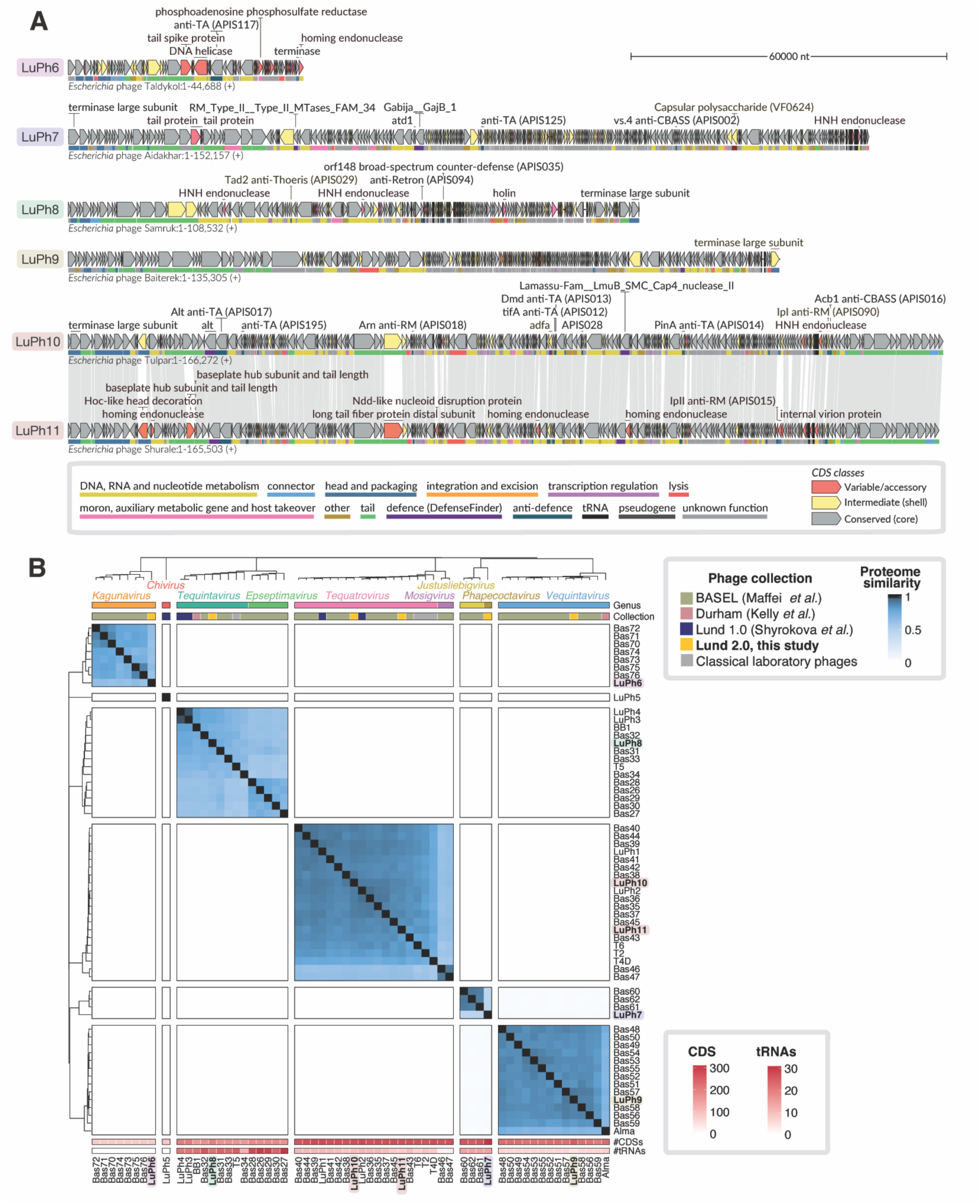
Genome organisation and of Lund Collection 2.0 phages and their relationships with other commonly used phages. **A)** Genome organisation of novel LuPh phages visualised using LoVis4u (Egorov & Atkinson, 2025). Gene fill colour corresponds to classes calculated based on the fraction of similar phage genomes in which they are encoded: conserved (core) genes are shown in grey, variable (cloud) genes in red, and intermediate (shell) genes in yellow. Functional annotations are indicated by coloured lines beneath each ORF, corresponding to the colour key at the bottom of the panel. A subset of non-hypothetical encoded proteins is labelled, including the large terminase subunit, proteins identified as variable by LoVis4u, and proteins with functional hits based on PyHMMER (Larralde & Zeller, 2023) annotation. **B)** Pairwise comparison of proteome composition similarity between phages from the Lund Phage Collection 2.0 (this study) and 1.0 (Shyrokova *et al*., 2025), extended BASEL (Humolli *et al*., 2025) and Durham (Kelly *et al*., 2023) collections as well as commonly used classical laboratory phages. Proteome composition similarity matrix values were calculated using LoVis4u (Egorov & Atkinson, 2025) as the fraction of shared homologous protein clusters between all pairs of genomes. The dendrogram is defined by hierarchical clustering of the proteome composition distance matrix.

Next, we tested the host range of the newly isolated LuPh phages using MG1655 ΔRM, MG1655 ΔRM *wbbL(+)*, BW25113 *uspC-flhDC*::IS5 and EV36 as hosts (**Figure 2**). Like the other members of *Kagunavirus* genus (Humolli *et al*., 2025), LuPh6 infection activity strictly depends on the O-antigen presence. Likewise, LuPh7, which was isolated using EV36 strain, can only infect EV36, but not the other strains we used. The *Tequintavirus* LuPh8 shows the broadest infectivity range. It infects MG1655 ΔRM irrespective of O-antigen presence, as well as BW25113, and EV36. Among the *Tequintavirus* representatives from BASEL collection, only Bas32 can infect O-antigen-positive *E. coli* (Maffei *et al*., 2021), and their ability to penetrate the K1 capsule has not been studied. We, therefore, tested all BASEL *Demerecviridae* phages ability to infect EV36. Like LuPh6, Bas32, but not the other phages, could infect EV36 cells (**Supplementary Figure 1A**). The *Vequintavirinae* isolate LuPh9 and the *Tequatrovirus* isolates LuPh10 and LuPh11 still can infect *E. coli* in the presence of O-antigen or the K1 capsule but show markedly smaller plaque size. While the ability of *Vequintavirinae* representatives to hydrolyse a variety of glycans is well-known (Kropinski *et al*, 2013; Maffei *et al*., 2021; Schwarzer *et al*, 2012), that of *Tequatrovirus* representatives is rather rare (Maffei et al., 2021). Finally, despite the strong similarity between LuPh8 and Salmonella phage vB_Sen_TUMS_E3 (Table 1), Luph8 could not infect *Salmonella enterica* subsp*. enterica* serovar Typhimurium (*Salmonella* Typhimurium) strain 12023s (**Supplementary Figure 1B**).

**Figure 2.**
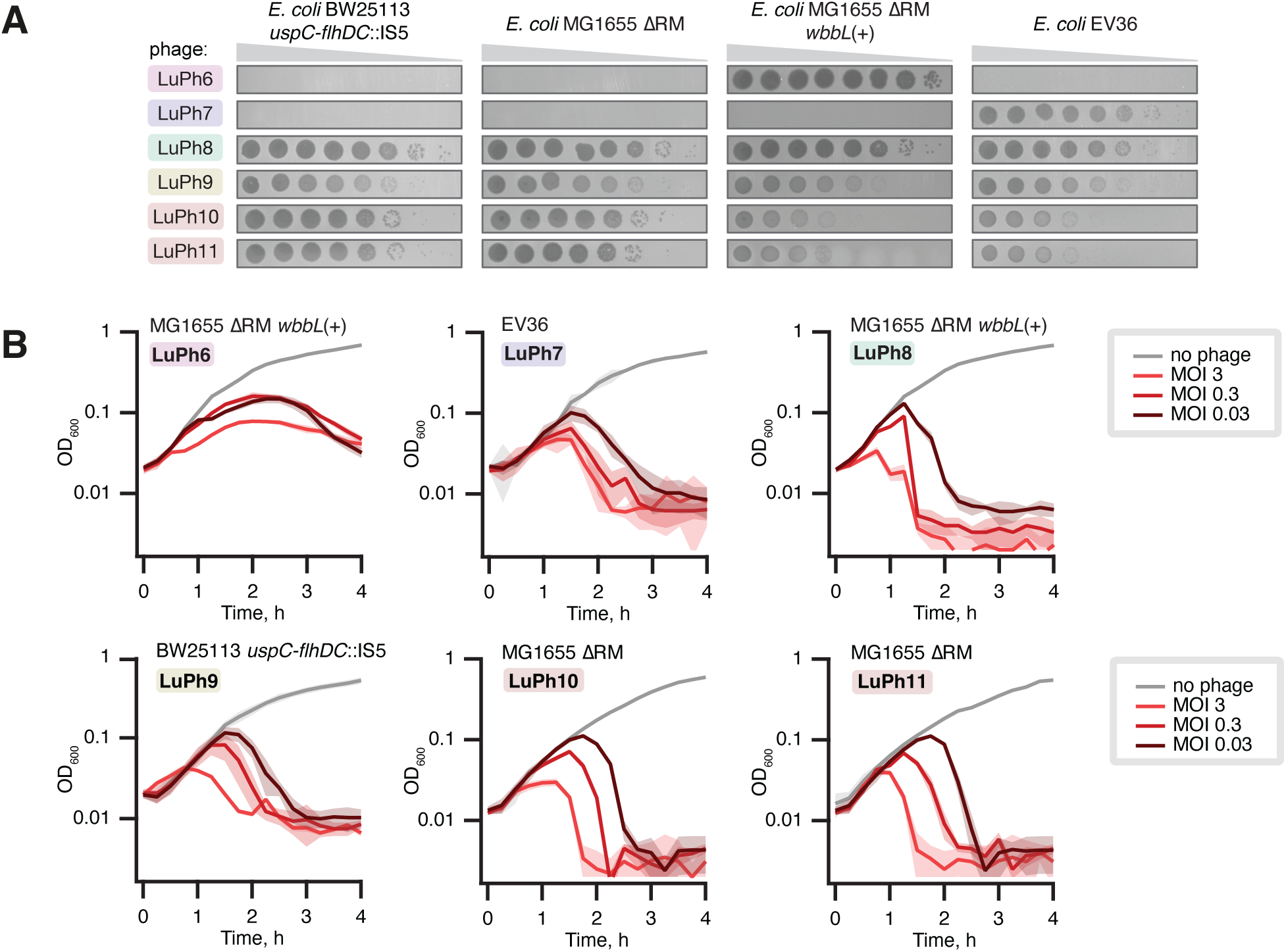
Host range analysis and infection kinetics of Lund Collection 2.0 phages. **A)** Serial dilution plaque assays on lawns of BW25113 *uspC*-*flhDC*::IS5 (VHB17), MG1655 ΔRM, MG1655 ΔRM *wbbL*(+), and EV36 *E. coli* cells. The LuPh phages were 10-fold serially diluted and spotted on the top agar plates followed by incubation at 37°C for 6 h. **B)** Growth curves of the indicated *E. coli* strains in the presence of the indicated phages at MOIs of 0, 0.03, 0.3, and 3. Each curve represents the average of three biological replicates, each using a different colony. Shaded areas indicate the standard deviation.

We also performed liquid culture phage infection assays to assess the dynamics of infection (**Figure 2B**). While all phages could lyse the appropriate *E. coli* strains, LuPh6 reproducibly showed no difference in the kinetics of infection between MOI 0.03 and 0.3.

## Materials and Methods

### Bacterial strains, plasmids and media

This study used four *E. coli* strains: two K-12 strains MG1655 ΔRM (F^−^ λ^−^ *ilvG^−^ rfb-50 rph-1* Δ*mrr-hsdRMS-mcrBC* Δ*mcrA)* and MG1655 ΔRM *wbbL(+)* where O16-type O-antigen expression is restored by deletion of the IS5 element that disrupted the *wbbL* gene (Maffei *et al*., 2021), BW25113 (F^−^ *λ^−^ Δ(araD-araB)567 Δ(rhaD-rhaB)568 ΔlacZ4787(::rrnB-3) hsdR514 rph-1 uspC-flhDC*::IS5) (VHB17) (Shyrokova *et al*., 2025) as well as a K12-K1 hybrid EV36 *rpsL9* [*argA*^+^ *kps*^+^ *rha*^+^]*^d^ galP23* (Vimr & Troy, 1985) and *Salmonella enterica* subsp*. enterica* serovar Typhimurium strain 12023s (ATCC 14028) (Humolli *et al*., 2025). *E. coli* strains were grown at 37°C in liquid or on solid (1.5% w/v agar) LB medium (10 g/L tryptone, 5 g/L yeast extract, and 10 g/L NaCl). When required for the pAS001 plasmid selection (Humolli *et al*., 2025), the medium was supplemented with 20 μg/mL chloramphenicol.

### Phage isolation, purification and preparation of phage stocks

Environmental water samples (50 mL each) were collected from Malyi Taldykol, an urban lake in Astana, Kazakhstan (51°07’00.7”N 71°23’13.5”E) on July 4, 2025. Samples were cleared from debris by centrifugation at 8,000 × g for 5 minutes at room temperature. Phages in the supernatant were precipitated by adding 1 mL of 2 M ZnCl_2_, followed by incubation at 37°C for 30 minutes. Following centrifugation at 8,000 × g for 10 minutes at room temperature, the pellet was resuspended in 0.5-1 mL of SM buffer (0.1 M NaCl, 0.01 M MgSO_4_, 0.05 M Tris-HCl, pH 7.5) and agitated at 600 rpm for 15 minutes at 37°C in a thermoblock. The suspension was then assayed for phages using the top agar overlay method (Kropinski *et al*., 2009). Briefly, 200 μL of the suspension was mixed with 200 μL of an overnight culture of *E. coli* BW25113 *uspC-flhDC*::IS5, EV36 or MG1655 ΔRM carrying pAS001 plasmid (Humolli *et al*., 2025) and incubated at room temperature for 15 minutes. The phage-cell mixture was then combined with 10 mL of top agar (0.5% w/v agar supplemented with 20 mM MgSO_4_ and 5 mM CaCl_2_) and poured onto square (12 × 12 cm) LB agar plates (1.5% w/v agar). Once the top agar had solidified, plates were incubated overnight at 37°C. Due to time constraints, individual plaques were re-streaked only once (using top agar overlays) during the course but were subsequently purified by at least three consecutive re-streaks.

Phage stocks were prepared by transferring the top agar from visible lysis zones of the final re-streak into 1 mL of SM buffer in a 2 mL tube. The mixture was vortexed for 30 seconds and then centrifuged at 14,000 rpm for 10 minutes at room temperature. The resulting supernatant (i.e., the phage stock) was transferred to a 1.5 mL tube and stored at 4 °C. For long-term storage, the phages were kept as virocells at −80°C (Golec *et al*, 2011).

### Liquid culture phage infection assays

Three independent overnight cultures of appropriate *E. coli* strains in LB medium supplemented with 10 mM MgSO_4_ and 2.5 mM CaCl_2_ were diluted to OD_600_=0.05 in the same medium, grown to OD600≈0.5, diluted with the same medium to OD_600_=0.075 and 100 μl were transferred to wells of a 96-well plate. Phage stocks were diluted in SM buffer to generate final MOI values of 3, 0.3, and 0.03 when 2.5 μl of the dilution was added to a well. Equal volume of SM buffer was added to the control wells (no phage). Bacterial growth was monitored at 37°C with double-orbital shaking at 400 rpm in a SPECTROstar Nano plate reader (BMG LABTECH) by measuring OD_600_ every 15 minutes for 6 hours.

### Efficiency of plating assays

Overnight cultures of *E. coli* were mixed with 10 ml top agar to a final concentration of 0.075 OD_600_ units/mL and overlaid onto LB agar plates (1.5% agar). Individual phage stocks were 10-fold serially diluted in SM buffer and 2.5 μL of each of the eight dilutions was spotted onto the solidified top agar. Plates were incubated at 37°C, and plaque formation was monitored at 6 and 24 hours.

### Phage genome sequencing, annotation and visualisation

Genomic DNA from bacteriophages was purified using the Phage DNA Isolation Kit (Norgen Biotek). Sequencing was performed by SeqVision company (Vilnius, Lithuania) with the following protocol: Bacteriophage gDNA was end-repaired with the addition of dA and adapters were ligated using a T/A ligation-based method without DNA fragmentation. The whole genome was amplified using barcoded primers annealing to the ligated adapters. The samples were purified, pooled, and nanopore native adapters were ligated (kit 14 chemistry). Samples were sequenced on R10.4.1 nanopore and basecalled using the super-accuracy v5.0.0 model on Dorado v0.9.5. Reads were demultiplexed using Dorado. The lowest quality 10% of reads were removed using Filtlong v0.2.1 (github.com/rrwick/Filtlong), the remaining reads were subsampled to 100-200 genome coverage and assembled using Flye v2.9.4 (Kolmogorov *et al*, 2019) to recover a circularized genome. The final assembly was polished using Medaka v2.0.1 (github.com/nanoporetech/medaka).

Phage assemblies were annotated using Pharokka v1.7.4 (Bouras *et al*., 2023). Specifically, coding sequences (CDSs) were predicted with PHANOTATE (McNair *et al*, 2019), tRNAs with tRNAscan-SE 2.0 (Chan *et al*, 2021), and tmRNAs with Aragorn (Laslett & Canback, 2004). Functional annotations of CDSs were assigned according to PHROGs categories (Terzian *et al*, 2021). The web version of BLASTn (Johnson *et al*., 2008) was used to query the novel phages against the NCBI nucleotide collection; summary tables of the results are provided in **Supplementary Table 1**.

To calculate pairwise nucleotide identity between the novel phages and their closest relatives identified by the NCBI search, we first used the DNAdiff v1.3 tool from the MUMmer4 aligner (Marcais *et al*, 2018) to determine the number of aligned nucleotides between genome pairs. To get intergenomic identity, the total number of aligned nucleotides from the query and reference genomes was then summed and normalised by the combined length of both genomes.

Genome organisation and proteome-level alignments were visualised using LoVis4u (Egorov & Atkinson, 2025), which employs MMseqs2 (Steinegger & Soding, 2017) for protein clustering and PyHMMER (Larralde & Zeller, 2023) searches against the DefenseFinder (Tesson *et al*, 2024) and dbAPIS (Yan *et al*, 2024) databases for additional functional annotation. LoVis4u was also used to construct a proteome composition similarity matrix, which was subsequently used for hierarchical clustering of phages and heatmap visualisation using ComplexHeatmap R package (Gu *et al*, 2016).

### Transmission electron microscopy

For negative staining, 4 µL of a filtered phage stock, appropriately diluted in SM buffer, was applied onto a carbon-coated 400-mesh copper grid (EMS) that had been glow-discharged in air for 60 seconds. The grids were blotted from the edge using Whatman grade 1 filter paper and negatively stained with 2% uranyl acetate for 60 seconds. The grids were imaged using a Thermo Scientific™ Talos™ L120C transmission electron microscope operating at 120 kV, which was equipped with a 4k × 4k Ceta CMOS camera. The dimensions of phage particles were measured using ImageJ (Schneider *et al*, 2012).

## Data availability

Phage genome sequences are available in the NCBI database under GenBank accession numbers PX880098-PX880103.

## Acknowledgements

We thank Fyodor A. Kondrashov and the organising committee of SMTB for organising the school, as well as the staff of Nazarbayev University for their support, Alexander Harms for sharing *E. coli* K-12 MG1655 ΔRM and Antonia Sagona for sharing *E. coli* EV36. This work was supported by the Carl Trygger Foundation (CTS24:3450 to M.J.O.J.), the Knut and Alice Wallenberg Foundation (2020-0037 to G.C.A. and V.H.), the Swedish Research Council (2022-01603, 2023-02353 and 2024-06071 to G.C.A.; 2021-01146 and 2024-06059 to V.H.), the Estonian Research Council (PRG2696 to V.H.), the Göran Gustafsson Foundation for Research in Natural Sciences and Medicine (the Göran Gustafsson Prize to V.H.), the Royal Physiographic Society of Lund (Endowments for the Natural Sciences, Medicine and Technology, number 45379 to A.A.E.). The computations were enabled by the Berzelius resource provided by the Knut and Alice Wallenberg Foundation at the National Supercomputer Centre, and by resources provided by the National Academic Infrastructure for Supercomputing in Sweden (NAISS), partially funded by the Swedish Research Council through grant agreement no. 2022-06725. The Cryo-EM for Life Sciences facility at Lund University and Lund University Bioimaging Centre (LBIC) are acknowledged for providing TEM support.

## References

Arredondo-Alonso S, Blundell-Hunter G, Fu Z, Gladstone RA, Fillol-Salom A, Loraine J, Cloutman-Green E, Johnsen PJ, Samuelsen O, Pontinen AK et al (2023) Evolutionary and functional history of the *Escherichia coli* K1 capsule. Nat Commun 14: 3294

Bouras G, Nepal R, Houtak G, Psaltis AJ, Wormald PJ, Vreugde S (2023) Pharokka: a fast scalable bacteriophage annotation tool. Bioinformatics 39

Chan PP, Lin BY, Mak AJ, Lowe TM (2021) tRNAscan-SE 2.0: improved detection and functional classification of transfer RNA genes. Nucleic Acids Res 49: 9077–9096

Cross AS, Gemski P, Sadoff JC, Orskov F, Orskov I (1984) The importance of the K1 capsule in invasive infections caused by *Escherichia coli*. J Infect Dis 149: 184–193

Datsenko KA, Wanner BL (2000) One-step inactivation of chromosomal genes in *Escherichia coli* K-12 using PCR products. Proc Natl Acad Sci U S A 97: 6640–6645

Dion MB, Oechslin F, Moineau S (2020) Phage diversity, genomics and phylogeny. Nat Rev Microbiol 18: 125–138

Egorov AA, Atkinson GC (2025) LoVis4u: a locus visualization tool for comparative genomics and coverage profiles. NAR Genom Bioinform 7: lqaf009

Golec P, Dabrowski K, Hejnowicz MS, Gozdek A, Los JM, Wegrzyn G, Lobocka MB, Los M (2011) A reliable method for storage of tailed phages. J Microbiol Methods 84: 486–489

Gong Q, Wang X, Huang H, Sun Y, Qian X, Xue F, Ren J, Dai J, Tang F (2021) Novel host recognition mechanism of the K1 capsule-specific phage of *Escherichia coli*: capsular polysaccharide as the first receptor and lipopolysaccharide as the secondary receptor. J Virol 95: e0092021

Grenier F, Matteau D, Baby V, Rodrigue S (2014) Complete Genome Sequence of *Escherichia coli* BW25113. Genome Announc 2

Gu Z, Eils R, Schlesner M (2016) Complex heatmaps reveal patterns and correlations in multidimensional genomic data. Bioinformatics 32: 2847–2849

Heller DM, Sivanathan V, Asai DJ, Hatfull GF (2024) SEA-PHAGES and SEA-GENES: Advancing Virology and Science Education. Annu Rev Virol 11: 1–20

Humolli D, Piel D, Maffei E, Heyer Y, Agustoni E, Shaidullina A, Willi L, Imwinkelried P, Estermann F, Cuenod A et al (2025) Completing the BASEL phage collection to unlock hidden diversity for systematic exploration of phage-host interactions. PLoS Biol 23: e3003063

Johnson M, Zaretskaya I, Raytselis Y, Merezhuk Y, McGinnis S, Madden TL (2008) NCBI BLAST: a better web interface. Nucleic Acids Res 36: W5–9

Kelly A, Went SC, Mariano G, Shaw LP, Picton DM, Duffner SJ, Coates I, Herdman-Grant R, Gordeeva J, Drobiazko A et al (2023) Diverse Durham collection phages demonstrate complex BREX defense responses. Appl Environ Microbiol 89: e0062323

King MR, Vimr RP, Steenbergen SM, Spanjaard L, Plunkett G, 3rd, Blattner FR, Vimr ER (2007) *Escherichia coli* K1-specific bacteriophage CUS-3 distribution and function in phase-variable capsular polysialic acid O acetylation. J Bacteriol 189: 6447–6456

Kolmogorov M, Yuan J, Lin Y, Pevzner PA (2019) Assembly of long, error-prone reads using repeat graphs. Nat Biotechnol 37: 540–546

Kropinski AM, Mazzocco A, Waddell TE, Lingohr E, Johnson RP (2009) Enumeration of bacteriophages by double agar overlay plaque assay. Methods Mol Biol 501: 69–76

Kropinski AM, Waddell T, Meng J, Franklin K, Ackermann HW, Ahmed R, Mazzocco A, Yates J, 3rd, Lingohr EJ, Johnson RP (2013) The host-range, genomics and proteomics of Escherichia coli O157:H7 bacteriophage rV5. Virol J 10: 76

Larralde M, Zeller G (2023) PyHMMER: a Python library binding to HMMER for efficient sequence analysis. Bioinformatics 39

Laslett D, Canback B (2004) ARAGORN, a program to detect tRNA genes and tmRNA genes in nucleotide sequences. Nucleic Acids Res 32: 11–16

Maffei E, Shaidullina A, Burkolter M, Heyer Y, Estermann F, Druelle V, Sauer P, Willi L, Michaelis S, Hilbi H et al (2021) Systematic exploration of Escherichia coli phage-host interactions with the BASEL phage collection. PLoS Biol 19: e3001424

Marcais G, Delcher AL, Phillippy AM, Coston R, Salzberg SL, Zimin A (2018) MUMmer4: A fast and versatile genome alignment system. PLoS Comput Biol 14: e1005944

McNair K, Zhou C, Dinsdale EA, Souza B, Edwards RA (2019) PHANOTATE: a novel approach to gene identification in phage genomes. Bioinformatics 35: 4537–4542

Rajab AAH, Fahmy EK, Esmaeel SE, Yousef N, Askoura M (2024) *In vitro* and *in vivo* assessment of the competence of a novel lytic phage vB_EcoS_UTEC10 targeting multidrug resistant *Escherichia coli* with a robust biofilm eradication activity. Microb Pathog 197: 107058

Schneider CA, Rasband WS, Eliceiri KW (2012) NIH Image to ImageJ: 25 years of image analysis. Nat Methods 9: 671–675

Scholl D, Merril C (2005) The genome of bacteriophage K1F, a T7-like phage that has acquired the ability to replicate on K1 strains of *Escherichia coli*. J Bacteriol 187: 8499–8503

Schwarzer D, Buettner FF, Browning C, Nazarov S, Rabsch W, Bethe A, Oberbeck A, Bowman VD, Stummeyer K, Muhlenhoff M et al (2012) A multivalent adsorption apparatus explains the broad host range of phage phi92: a comprehensive genomic and structural analysis. J Virol 86: 10384–10398

Shyrokova L, Egorov AA, Cole A, Duque-Pedraza JJ, Tyagi A, Ernits K, Mets T, Kurata T, Juozapaitis J, Yang ALJ et al (2025) Characterization of five environmental phages infecting Escherichia coli K-12 isolated during a phage biology training course. Microbiol Spectr: e0227425

Steinegger M, Soding J (2017) MMseqs2 enables sensitive protein sequence searching for the analysis of massive data sets. Nat Biotechnol 35: 1026–1028

Terzian P, Olo Ndela E, Galiez C, Lossouarn J, Perez Bucio RE, Mom R, Toussaint A, Petit MA, Enault F (2021) PHROG: families of prokaryotic virus proteins clustered using remote homology. NAR Genom Bioinform 3: lqab067

Tesson F, Planel R, Egorov AA, Georjon H, Vaysset H, Brancotte B, Néron B, Mordret E, Atkinson GC, Bernheim A et al (2024) A Comprehensive Resource for Exploring Antiphage Defense: DefenseFinder Webservice,Wiki and Databases. Peer Community Journal 4

Turner D, Kropinski AM, Adriaenssens EM (2021) A Roadmap for Genome-Based Phage Taxonomy. Viruses 13

Turner D, Shkoporov AN, Lood C, Millard AD, Dutilh BE, Alfenas-Zerbini P, van Zyl LJ, Aziz RK, Oksanen HM, Poranen MM et al (2023) Abolishment of morphology-based taxa and change to binomial species names: 2022 taxonomy update of the ICTV bacterial viruses subcommittee. Arch Virol 168: 74

Vimr ER, Troy FA (1985) Identification of an inducible catabolic system for sialic acids (*nan*) in *Escherichia coli*. J Bacteriol 164: 845–853

Yan Y, Zheng J, Zhang X, Yin Y (2024) dbAPIS: a database of anti-prokaryotic immune system genes. Nucleic Acids Res 52: D419–D425

