## Supplementary Figure 1 for "Characterization of six environmental *coli*-phages isolated in Astana, Kazakhstan, during the School of Molecular and Theoretical Biology"

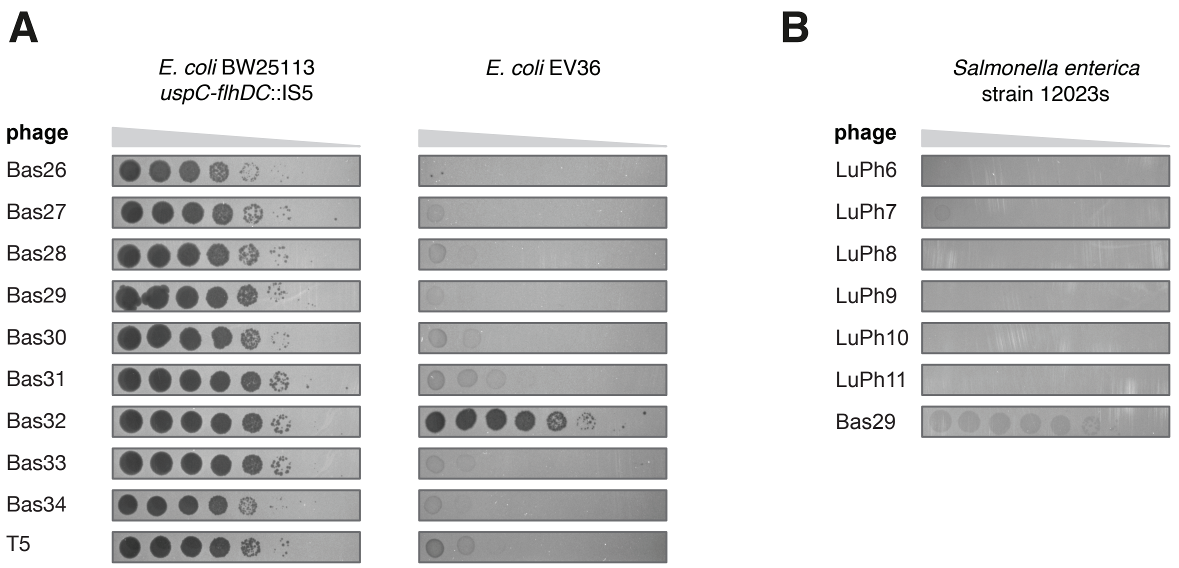


**Supplementary Figure 1**. Host range analysis of BASEL and Lund Collection 2.0 phages. A) Serial dilution plaque assays on lawns of BW25113 uspC-flhDC::IS5 (VHB17) and EV36 *E. coli* cells. The BASEL representatives of *Demerecviridae* phages were 10-fold serially diluted and spotted on the top agar plates followed by incubation at 37°C for 6 h. B) Serial dilution plaque assays on a lawn of *Salmonella enterica* strain 12023s. The Lund Collection 2.0 phages were 10-fold serially diluted and spotted on the top agar plates followed by incubation at 37°C for 6 h.
