## Supplementary figures and images for "Characterization of six environmental *coli*-phages isolated in Astana, Kazakhstan, during the School of Molecular and Theoretical Biology"

### Supplementary File 3

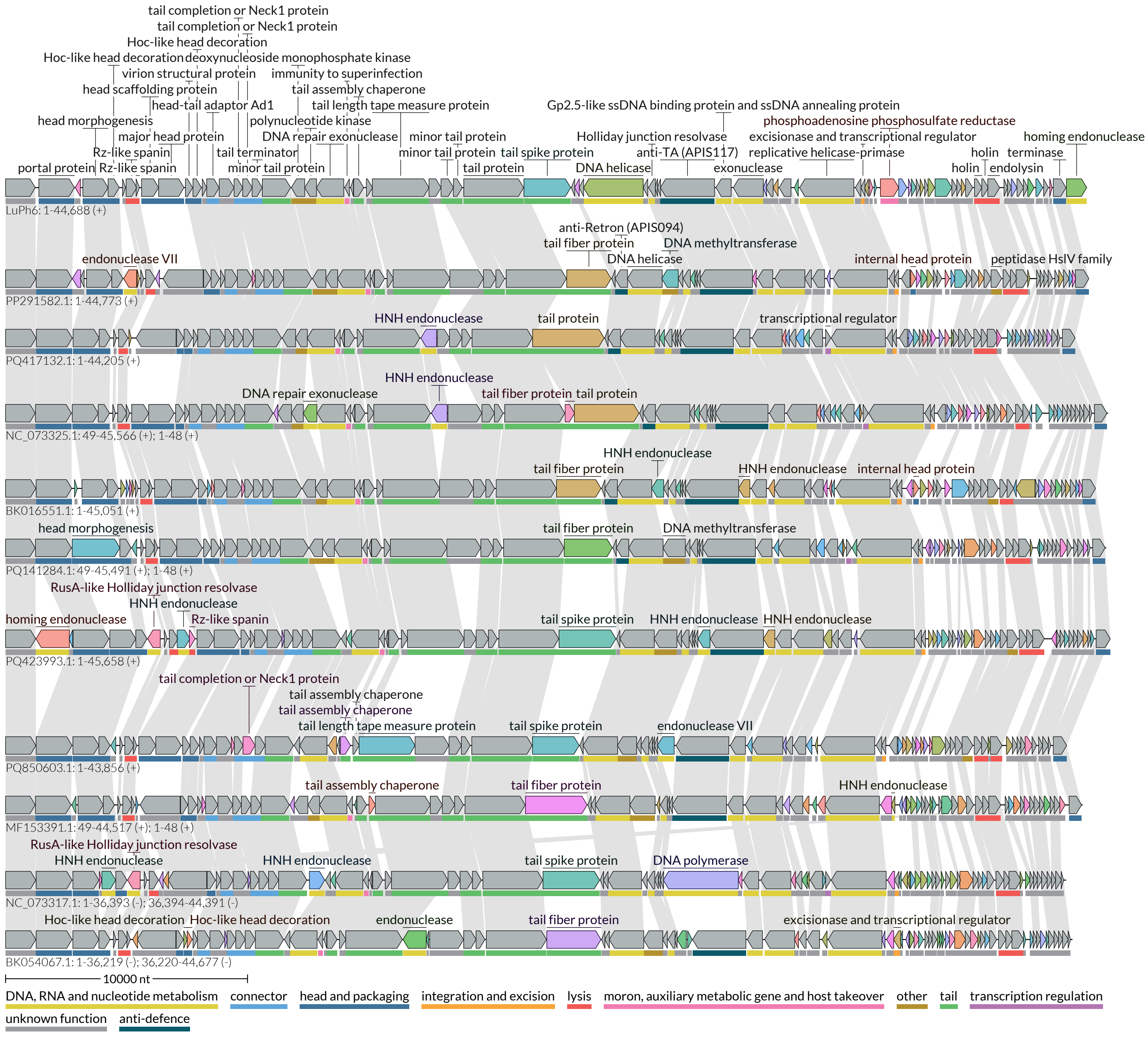

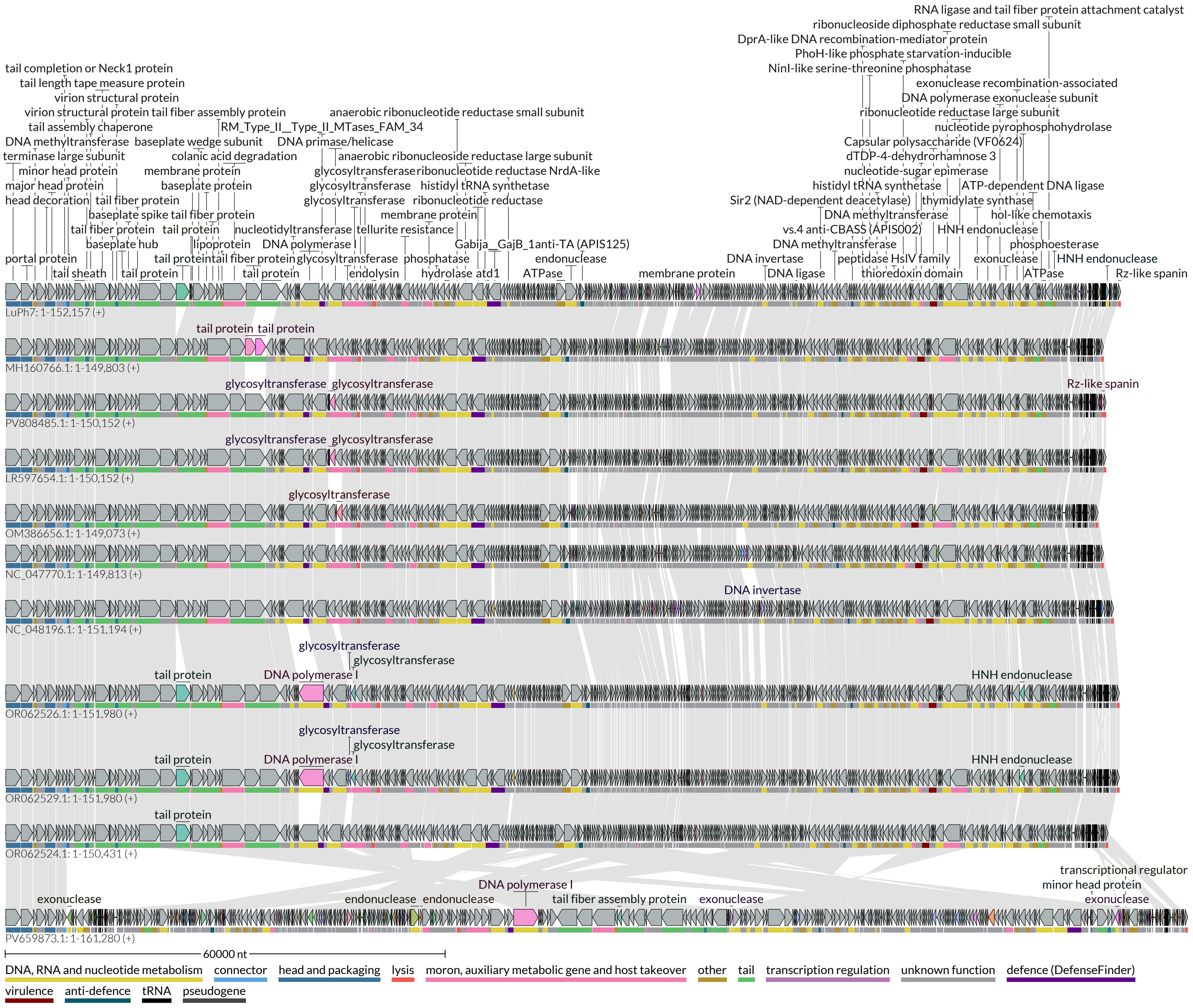

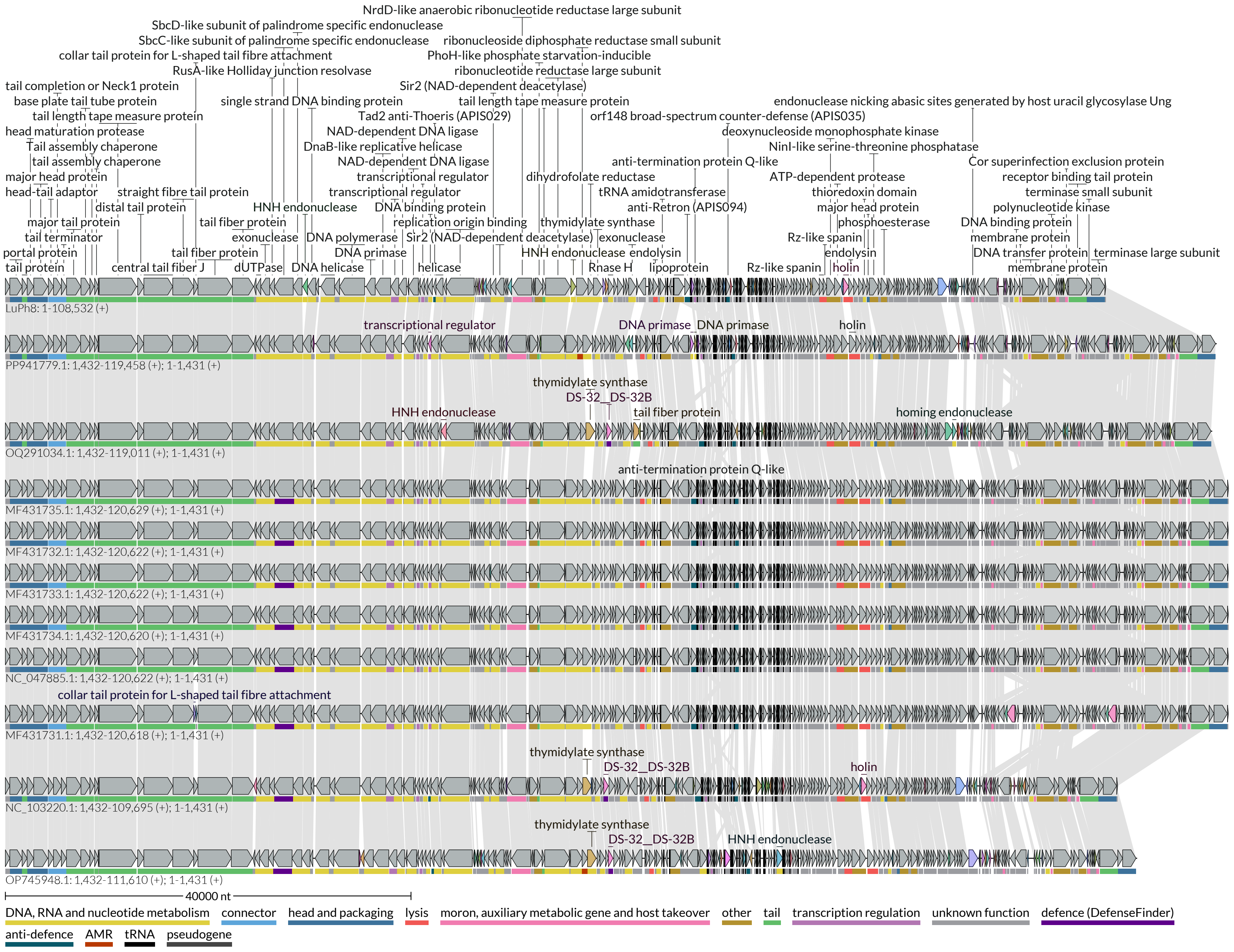

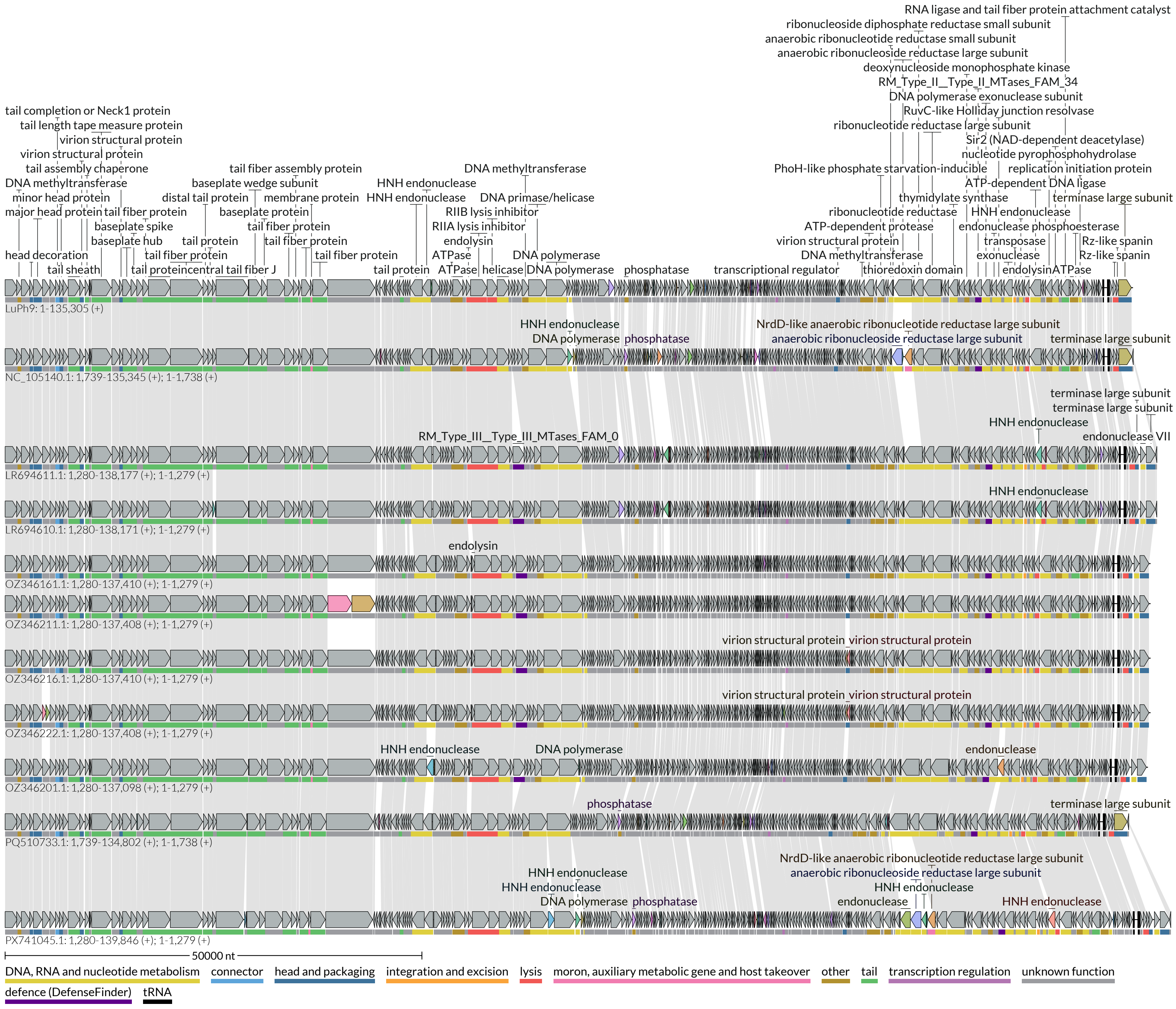

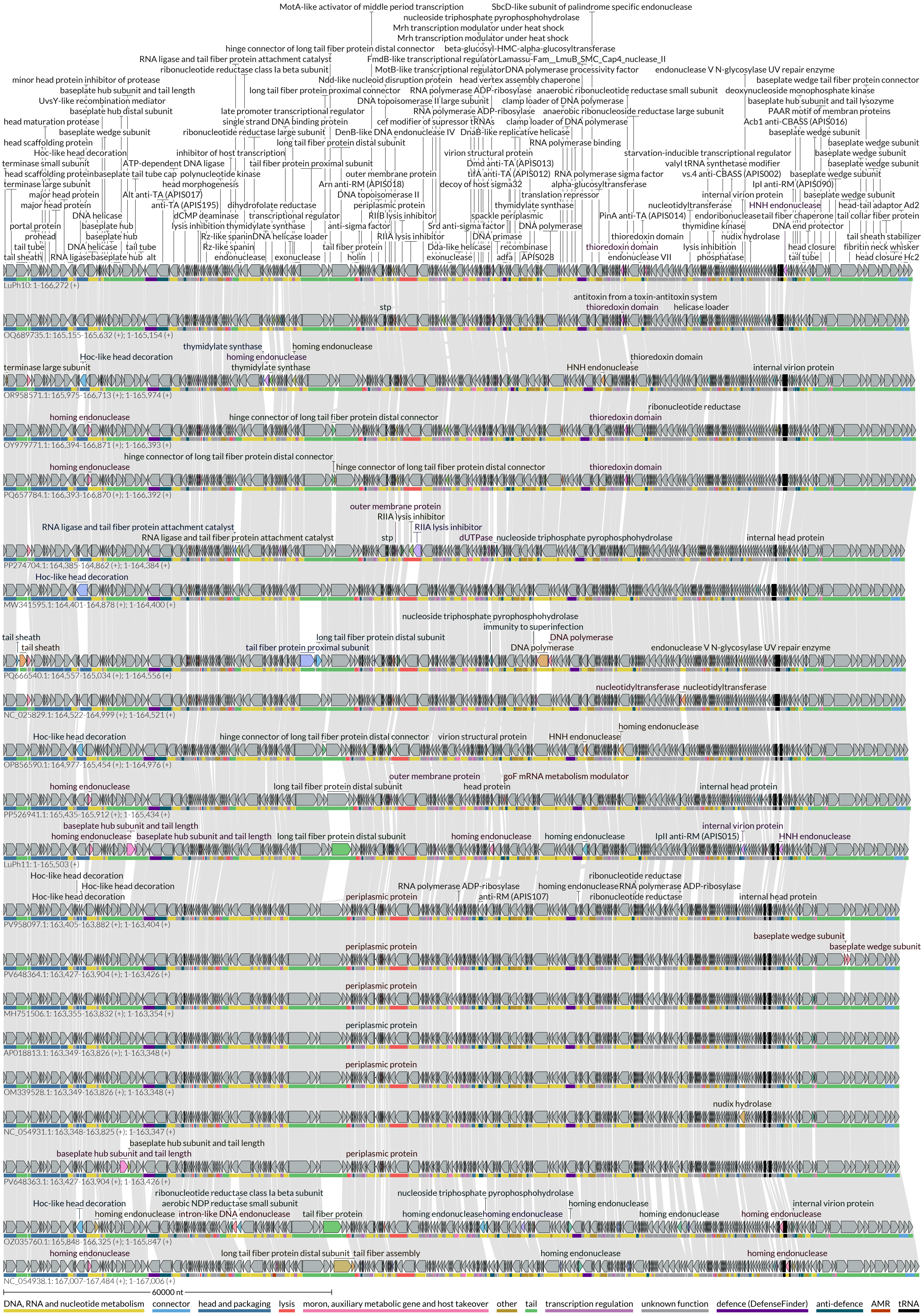

### Supplementary File 4

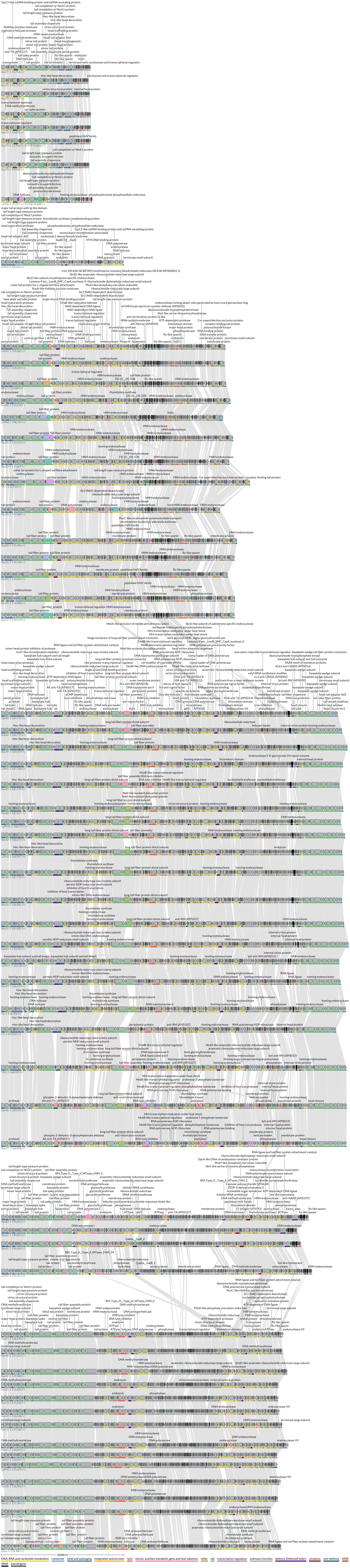
